# PTEN deficiency linked to chromosome 10q loss leads to aggressive NF2 mutant meningioma biology

**DOI:** 10.64898/2025.12.21.695820

**Authors:** Abigail G. Parrish, H. Nayanga Thirimanne, Sonali Arora, Christel Herold-Mende, Philipp Sievers, Felix Sahm, Frank Szulzewsky, Eric C. Holland

## Abstract

While most meningiomas are benign and can be surgically removed, a subset behaves aggressively, recurs quickly, and can ultimately be fatal. Recent work has focused on defining this aggressive group. To better characterize this clinically distinct, high-risk group, we analyzed bulk RNA sequencing (RNA-seq) from human meningiomas using a dimension-reduced reference landscape. We identified an NF2 mutant subtype enriched for chromosome 10q loss and low PTEN expression, both of which were strongly associated with shorter time to recurrence. To directly test causality, we engineered several genetically accurate, immunocompetent *in vitro* and *in vivo* mouse models with *Pten* knockdown or knockout. Across models, *Pten* loss markedly accelerated YAP1-driven tumorigenesis. Projection of *Pten*-deficient mouse tumors onto bulk and single-cell RNA-seq datasets from human meningiomas revealed that mouse tumors faithfully recapitulate the transcriptional programs of aggressive NF2 mutant meningiomas and robustly predicted clinical outcomes. Collectively, these findings demonstrate that PTEN loss cooperates with YAP1 activation to drive aggressive NF2 mutant meningioma biology, and that the high fidelity of this model to human disease establishes it as a robust platform for mechanistic investigation and therapeutic testing. More broadly, this work highlights the power of cross-species integration in both validating preclinical models and enhancing the translational relevance and utility of dimension-reduced reference landscapes across diverse disease types.

**One Sentence Summary:** PTEN loss causally drives aggressive NF2 mutant meningiomas, and cross-species integration provides a novel, high-fidelity system to validate preclinical models to human disease.

## INTRODUCTION

Meningiomas are the most common intracranial neoplasms, accounting for approximately 40% of all brain tumors, with a marked enrichment in adults aged 65 years and older (*1*). Although more than 80% of meningiomas are classified as grade 1 according to the fifth edition of the 2021 WHO Classification of Tumors of the Central Nervous System (CNS5), a subset exhibits aggressive, life-threatening behavior irrespective of histologic grade (*1*).

Nearly half of all meningiomas harbor a loss of one copy of chromosome 22q, which contains the NF2 tumor suppressor gene, the most frequent genetic aberration identified in these tumors (*2, 3*). NF2 is the gene that encodes the protein Merlin, a key upstream regulator of the Hippo signaling pathway that governs organ size and translates mechanical stimuli into transcriptional programs. When Hippo signaling is active, the transcriptional coactivators YAP1 and its paralog TAZ are phosphorylated through a core kinase cascade, leading to nuclear exclusion, cytoplasmic sequestration, and proteasomal degradation. Loss of NF2 function, a central driver in several nervous system tumors such as ependymomas, mesotheliomas, and vestibular schwannomas, decreases the activity of the pathway, thereby increasing and deregulating the activity of YAP1 (*4*).

Previous studies have demonstrated that a constitutively active YAP1 mutant, which is characterized by a combined S127/397A mutation (2SA-YAP1), can escape Hippo pathway regulation, deregulating and activating oncogenic YAP1. This constitutively active YAP1 mimics NF2 loss and is both necessary and sufficient to drive YAP1-mediated meningioma formation in mice (*4, 5*). While high-grade and recurrent tumors arise across all molecular subtypes, they are notably enriched among NF2 mutant meningiomas. Benign NF2 mutant meningiomas typically exhibit chromosome 22q loss with few additional genomic changes, whereas aggressive NF2 mutant tumors display complex genomic landscapes characterized by the additional recurrent losses of chromosomes 1p, 14q, 10q, 9p, and 18q, as well as gains of chromosomes 1q, 17q, and 20q (*6, 7*). Mutations in TERT promoter, CDKN2A, ARID1A, PTEN, KDM6A, TP53, and SUFU are among the most frequently altered genes in high-grade meningiomas (*8, 9*).

We have recently generated a comprehensive bulk RNA-seq-based reference landscape using Uniform Manifold Approximation and Projection (UMAP) of 1,298 human tumors, clustering samples by global gene expression patterns (*10*). Tumors with functional NF2 inactivation and chromosome 22q loss formed a dominant cluster, where high-grade tumors (enriched for WHO CNS5 grades 2-3) concentrated toward one end, suggesting a shared, distinct transcriptomic profile. Notably, low-grade tumors (i.e. grade 1) within this region showed recurrence times comparable to high-grade tumors, suggesting that histologic grade alone does not fully capture tumor aggressiveness. Instead, since clustering is based on global gene expression, these findings suggest that transcriptomic signatures more accurately predict clinical behavior, with reduced recurrence times likely driven by specific gene expression programs associated with tumor progression (*11, 12*).

Within the aggressive NF2 mutant region of the UMAP, tumors also exhibited frequent losses of chromosomes 1p, 6q, 10q, and 14q. Patients with these copy number alterations (CNAs) tend to have shorter recurrence intervals compared to those without such changes (*10, 13–15*). The loss of chromosome 10q is hypothesized to drive tumor progression through deletion of the PTEN tumor suppressor gene (harbored on chromosome 10q), promoting oncogenic signaling and more aggressive phenotypes. PTEN is a critic al negative regulator of the PI3K/AKT/mTOR/S6 signal transduction pathway, which governs cell growth, survival, and proliferation.

Loss of PTEN has been implicated in several malignancies, including glioblastoma, breast, endometrial, and prostate cancers, and has also been reported in meningiomas (*13, 16–18*). While loss of chromosome 10q and PTEN alterations are uncommon in low-grade meningiomas, it may contribute to malignant progression in anaplastic forms (*17*). There is a strong association between PTEN-related Cowden Syndrome and meningioma occurrence (*18*). Additionally, PTEN mutations co-occurring with NF2 loss and combined NF2 and PTEN mutations correlate with higher WHO grades, increased proliferation, recurrence, and mortality (*13, 16*). Despite these insights, the mechanistic contribution of PTEN loss to meningioma pathogenesis remains poorly understood.

In this study, we leveraged our meningioma UMAP to analyze the transcriptomic landscape of meningiomas and interrogate the biology underlying aggressive NF2 mutant tumors. Our analyses reveal that these tumors exhibit regionally aggressive behavior associated with WHO grade and chromosomal alterations, particularly loss of chromosome 10q encompassing the PTEN locus. *Pten* deficiency enhances YAP1-driven cell growth *in vitro,* and mouse models demonstrate that concurrent YAP1 activation and *Pten* loss increase tumor aggressiveness (both in the presence or absence of additional *Cdkn2a* loss). Cross-species projection of *Pten*-deficient mouse tumors onto bulk and single-cell RNA-seq datasets from human meningiomas demonstrated that these models mimic the core biological signatures of highly aggressive NF2 mutant meningiomas and accurately stratify clinical outcomes. These data establish cooperative YAP1 activation and PTEN loss as a central driver of highly aggressive NF2 mutant meningioma biology and introduce cross-species projection as a generalizable framework for aligning preclinical models with the human disease states they are designed to mimic, with broad implications for translational disease research.

## RESULTS

### UMAP reveals a distinct aggressive NF2-mutant meningioma state defined by chromosome 22q and CDKN2A loss

We have previously generated a transcriptomic landscape of 1,298 bulk RNA-seq profiles from human meningiomas by applying UMAP to batch-corrected, normalized transcript counts (*10*). Global gene expression delineated meningioma subtypes, which were corroborated by clinical and genomic metadata, and independently validated using DBSCAN clustering.

Bulk RNA-seq analysis revealed clear transcriptional stratification of meningioma samples in functional clusters that correlated with WHO CNS5 grade, with grade 3 tumors with the shortest time to recurrence, followed by grade 2 and 1, respectively (Fig. 1A-B). Clusters also distinguish NF2 status and tumor aggressiveness, where aggressive NF2 mutant tumors localized to one end of the NF2 mutant region, whereas benign NF2 mutant tumors occupied the opposite end. NF2 wild-type tumors formed a distinct cluster on the opposite side of the UMAP (Fig. 1C). Kaplan-Meier analysis showed the shortest time to recurrence in aggressive NF2 mutant tumors, intermediate recurrence in NF2 wild-type tumors, and the longest recurrence-free rate in benign NF2 mutant tumors (Fig. 1D).

**Fig. 1.**
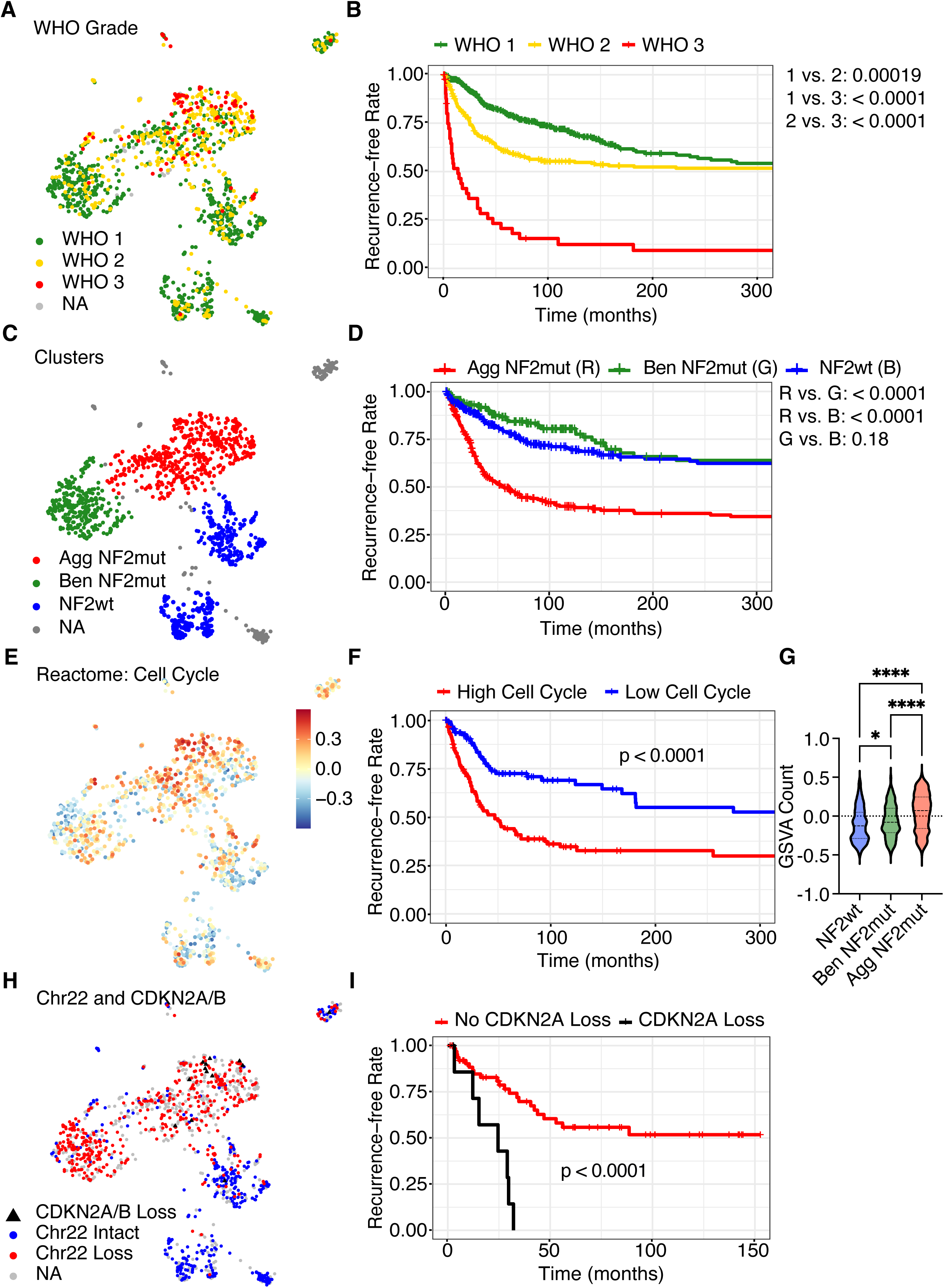
UMAP reveals a distinct aggressive NF2-mutant meningioma state defined by chromosome 22q and CDKN2A loss. UMAP colored by WHO grade and its corresponding Kaplan-Meier curve (**A–B**). UMAP colored by clusters and its corresponding Kaplan-Meier curve (**C–D**). UMAP colored by cell cycle GSVA and its corresponding Kaplan-Meier curve (**E–F**). Violin plot of cell cycle GSVA scores across clusters where NF2wt vs. Ben NF2mut p = 0.0369, NF2wt vs. Agg NF2mut p < 0.0001, and Ben NF2mut vs. Agg NF2mut p < 0.0001 (**G**). UMAP colored by Chromosome 22q and CDKN2A/B status and its corresponding Kaplan-Meier curve (**H-I**). Statistical analyses for violin plots were performed using one-way ANOVA, and Kaplan–Meier survival analyses were conducted using the log-rank (Mantel–Cox) test. Significance thresholds were defined as p > 0.05 (ns), p ≤ 0.05 (*), p ≤ 0.01 **(**)**, p ≤ 0.001 (***), and p ≤ 0.0001 (****), unless otherwise indicated.

Aggressive NF2 mutant tumors were markedly enriched for elevated cell-cycle activity, reflected by significantly higher cell-cycle gene set variation analysis (GSVA) scores (Fig. 1E). Stratification of patients by the upper and lower quartiles of cell-cycle GSVA values demonstrated that heightened cell-cycle gene expression was associated with significantly shorter recurrence intervals (Fig. 1F). Aggressive NF2 mutant tumors had significantly higher GSVA scores than benign NF2 mutant and NF2 wild-type tumors (Fig. 1G).

Genomic alterations mirrored these transcriptional features. Consistent with previous studies, concurrent loss of NF2 on chromosome 22q and CDKN2A was enriched in aggressive NF2 mutant region of the UMAP (Fig. 1H). Patients with CDKN2A loss showed substantially shorter recurrence times relative to those without this alteration (Fig. 1I).

### Aggressive NF2-mutant meningiomas are driven by recurrent chromosomal aneuploidies, including chromosome 10q loss encompassing PTEN

Aggressive NF2 mutant meningiomas exhibited extensive copy number alterations beyond chromosome 22q loss, notably, loss of chromosome 10q, which harbors the PTEN tumor suppressor locus (Fig. 2A). Among 132 patients with chromosome 10q loss across the 1,298 total samples across the UMAP, 105 (approximately 80 percent) localized to this region. This accounts for nearly a quarter of patients within the aggressive NF2 mutant region (476 total). Clinically, chromosome 10q loss was associated with shorter times to recurrence within aggressive NF2 mutant tumors (Fig. 2B). Consistent with this alteration, PTEN expression was significantly reduced in the aggressive NF2 mutant region compared to the benign NF2 mutant and NF2 wild-type tumors (Fig. 2C-D). Combining both NF2 mutant clusters (benign and aggressive) revealed that samples harboring chromosome 10q loss exhibited significantly lower PTEN expression levels, and this lower expression of PTEN correlated with shorter time to recurrence (Fig. 2E-F). Within the aggressive NF2 mutant cluster, tumors harboring chromosome 10q loss again showed significantly reduced PTEN expression, with this downregulation associated with a nonsignificant trend toward shorter time to recurrence (Fig. 2G-H). Low PTEN levels predicted poor outcomes even in some tumors lacking detectable chromosome 10q loss (Suppl. Fig. S2A-B).

**Fig. 2.**
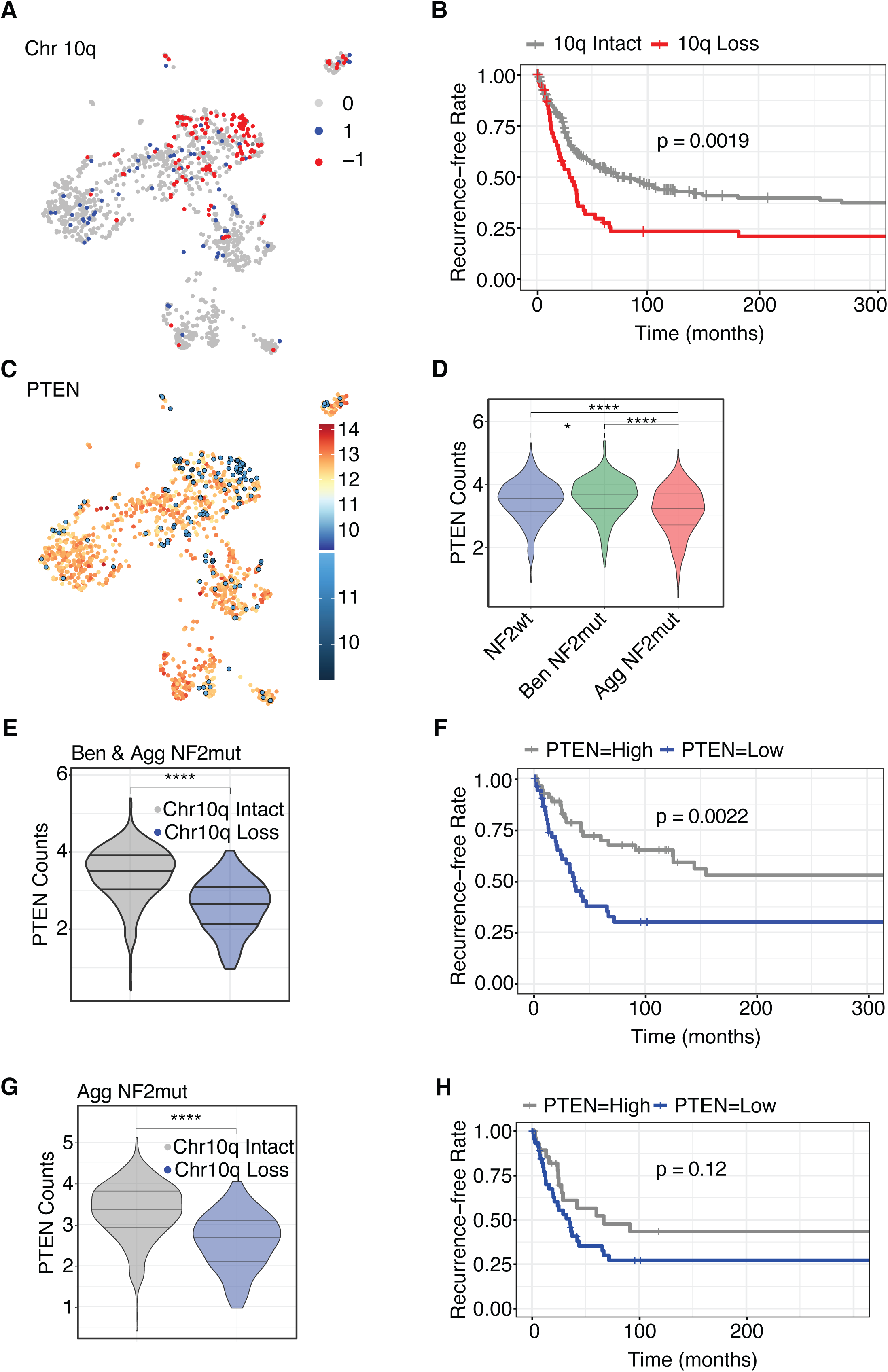
Aggressive NF2-mutant meningiomas are driven by recurrent chromosomal aneuploidies, most prominently chromosome 10q loss encompassing PTEN. UMAP colored by chromosome 10q status and its corresponding Kaplan-Meier curve (**A–B**). UMAP colored by PTEN RNA expression (**C**). Violin plot of PTEN expression across clusters, where NF2wt vs. Ben NF2mut p = 0.022, NF2wt vs. Agg. NF2mut p = 7E-11, and Ben NF2mut vs. Agg NF2mut p = 8.6E-16 (**D**). Violin plot of PTEN expression and corresponding Kaplan-Meier in chromosome 10q-loss versus 10q-wild-type tumors within NF2 mutant regions where p = 2.22E-16 (**E–F**). Violin plot of PTEN expression and corresponding Kaplan-Meier in chromosome 10q-loss versus 10q-wild-type tumors within the aggressive NF2 mutant region where p = 2.22E-16 (**G–H**). Statistical analyses for violin plots were performed using one-way ANOVA, and Kaplan–Meier survival analyses were conducted using the log-rank (Mantel–Cox) test. Significance thresholds were defined as p > 0.05 (ns), p ≤ 0.05 (*), p ≤ 0.01 (**), p ≤ 0.001 (***), and p ≤ 0.0001 (****), unless otherwise indicated.

### RCAS/tv-a mouse models of YAP1-driven meningiomas are more aggressive with concurrent loss of *Cdkn2a* and *Pten*

To explore the effects of PTEN loss in a preclinical context, we generated primary arachnoid cap cell cultures from neonatal *Nestin* (*N*)/tv-a; *Cdkn2a*^−/−^; *Pten*^fl/fl^ mice. Cells were transduced with replication-competent DF-1 fibroblast supernatant expressing constitutively active YAP1 (S127/397A-(2SA)-YAP1) either alone or together with Cre recombinase (deleting *Pten*). Arachnoid cells expressing both 2SA-YAP1 and Cre exhibited lower expression of PTEN and elevated phosphorylation of AKT and S6 compared to 2SA-YAP1-only cells, suggesting PI3K pathway activation upon Cre-mediated *Pten* knockout (Fig. 3A). 2SA-YAP1/Pten-null arachnoid cells proliferated more rapidly than 2SA-YAP1/*Pten*-wild-type cells (Fig. 3B). Consistent with these findings, subarachnoid injection of *Pten*-null arachnoid cells into neonatal and adult mice resulted in meningioma-like tumor formation in 100% of animals within 50 days, whereas 2SA-YAP1/*Pten*-wild-type cells failed to form tumors (Fig. 3C). Histology and magnetic resonance imaging shows that these tumors frequently grew outside the brain’s parenchyma and formed large intracranial (IC), extra-axial (EA), and extracranial (EC) tumors (Suppl. Fig. S3A).

**Fig. 3.**
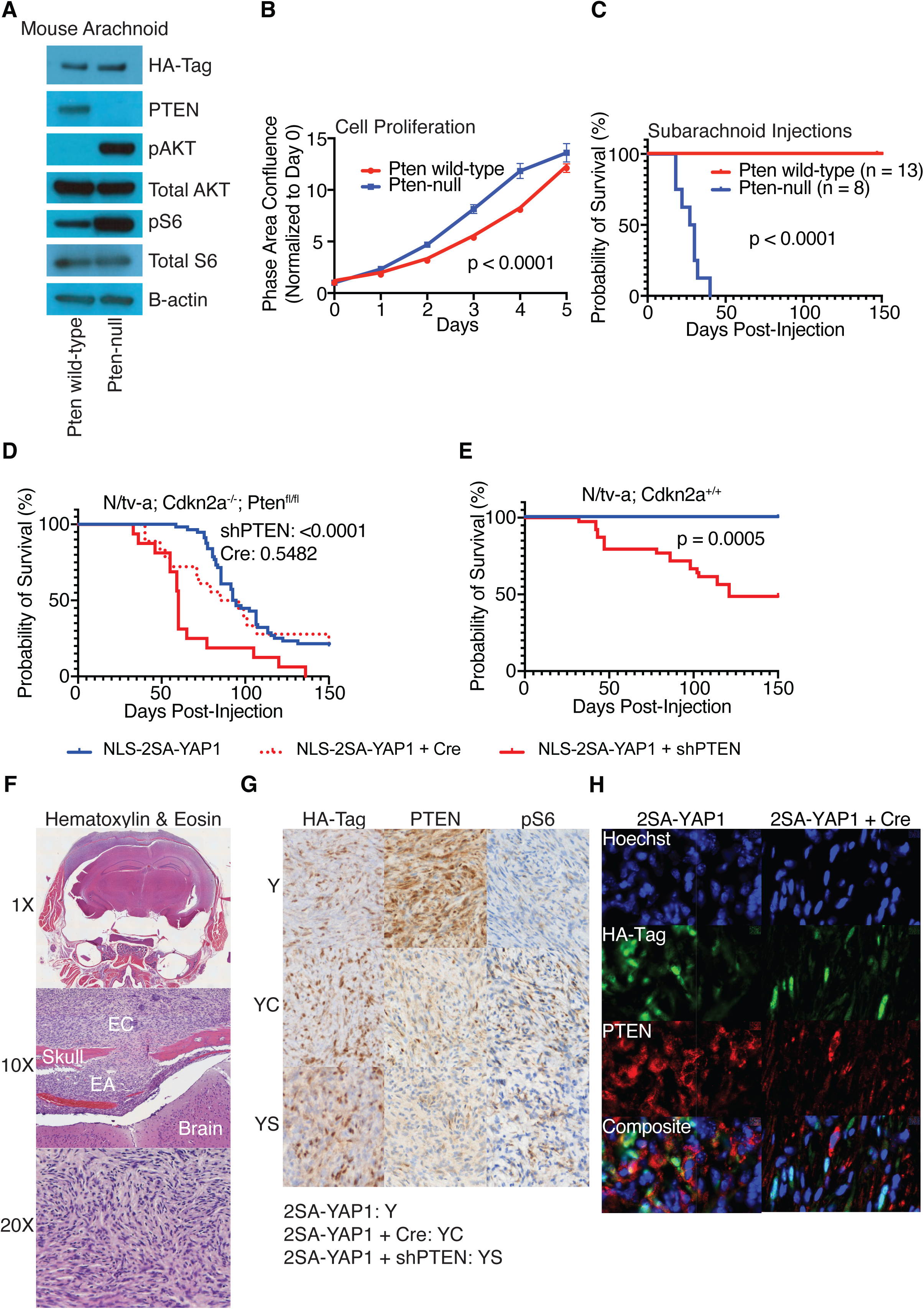
RCAS/tv-a mouse models of YAP1-driven meningiomas are more aggressive with concurrent loss of *Cdkn2a* and *Pten*. Western blot and proliferation assay of primary arachnoid cap cells from (*N*)/tv-a; *Cdkn2a*^−/−^; *Pten*^fl/fl^ mice transduced with YAP1 with or without Cre (**A–B**). Subarachnoid implantation of arachnoid cells expressing YAP1 with or without Pten loss (**C**). *In vivo* RCAS/tv-a–mediated expression of 2SA-YAP1 with or without *Pten* knockdown or deletion in neonatal mice (**D–E**). Hematoxylin & eosin of *Pten*-deficient mouse tumor, with extra-axial (EA) and extra-cranial (EC) tumors shown (**F**). Immunohistochemistry of mouse tumors for HA-tag, PTEN, and phospho-S6 (**G**). Immunofluorescence for PTEN status in HA-positive tumor cells (**H**). Statistical analyses were conducted using the log-rank (Mantel–Cox) test. Significance thresholds were defined as p > 0.05 (ns), p ≤ 0.05 (*), p ≤ 0.01 **(**)**, p ≤ 0.001 (***), and p ≤ 0.0001 (****), unless otherwise indicated.

Previous studies have shown that the RCAS-mediated expression of 2SA-YAP1 can drive the formation of tumors that resemble NF2 mutant meningiomas in *Cdkn2a* null neonatal mice (*4*). To further assess whether *Pten* loss functionally cooperates with YAP activation *in vivo*, we employed the RCAS/tv-a system for postnatal somatic gene transfer to express constitutively active YAP1 in the subarachnoid space of neonatal (*N*)/tv-a; *Cdkn2a*^−/−^; *Pten*^fl/fl^ mice. Injection of 2SA-YAP1 into Cdkn2a-null mice induced meningiomas in the subarachnoid space as previously observed (*4*). Additional *Pten* loss via shRNA-mediated knockdown accelerated tumor formation compared to 2SA-YAP1 expression alone (Fig. 3D). Of note, Cre-mediated complete deletion of *Pten* only led to a non-significant trend of accelerated tumor growth (Fig. 3D). RNA-seq confirmed that reads aligned to the Cre gene, suggesting that *Pten* was successfully excised from the *Pten* knockout tumors (Suppl. Fig. S3B).

We then explored whether *Pten* loss can also accelerate 2SA-YAP1-mediated tumor formation in a *Cdkn2a* wild-type background. We injected 2SA-YAP1 alone or co-injected 2SA-YAP1 and a short hairpin RNA (shRNA) against *Pten* into (*N*)/tv-a; *Cdkn2a* wild-type mice. shRNA-mediated *Pten* knockdown also significantly accelerated tumor formation in *Cdkn2a* wild-type background. In *Cdkn2a* wild-type mice, *Pten* knockdown was required for tumor formation, as mice receiving 2SA-YAP1 alone did not develop tumors (Fig. 3E, Suppl. Fig. S3C).

Experimental RCAS/tv-a mouse tumors shared histomorphological features with human meningiomas, characterized by compact spindled cells and fibrous whorls, and frequently grew extra-axially (EA) and extracranially (EC) (Fig. 3F). Additionally, tumors were positive for HA-tag staining, confirming that they were driven by the constitutively active YAP1 transgene (Fig. 3G). Compared with tumors expressing YAP1 alone, *Pten*-deficient tumors displayed reduced PTEN immunoreactivity and increased phospho-S6 staining, indicating robust activation of the PI3K pathway. Immunofluorescence showed that PTEN was absent in HA-positive tumor cells, with residual *Pten* detectable only in non-tumor cells (Fig. 3H).

### Bulk RNA-seq from mouse models distinguishes *Pten* status *in vivo*

We isolated total RNA from tumors driven by 2SA-YAP1 alone, and those driven by both 2SA-YAP1 and Cre- or shRNA-mediated *Pten* loss in *(N)*/tv-a; *Cdkn2a*^−/−^; *Pten*^fl/fl^ mice. After performing RNA sequencing (RNA-seq), principal component analysis (PCA) and multidimensional scaling clearly separated tumors driven by 2SA-YAP1 alone, and those driven by both 2SA-YAP1 and Cre- or shRNA-mediated *Pten* loss (Fig. 4A, Suppl. Fig. S4A). Heatmap analysis revealed unique transcriptional patterns in *Pten*-deficient models (Fig. 4B). We calculated differentially expressed genes between *Pten*-deficient tumors and those driven by 2SA-YAP1 alone (Suppl. Table S4A–C). We observed significant changes in gene expression in *Pten*-deficient samples, identifying a core signature of 451 up-regulated DEGs and 521 down-regulated DEGs (Fig. 4C).

**Fig. 4.**
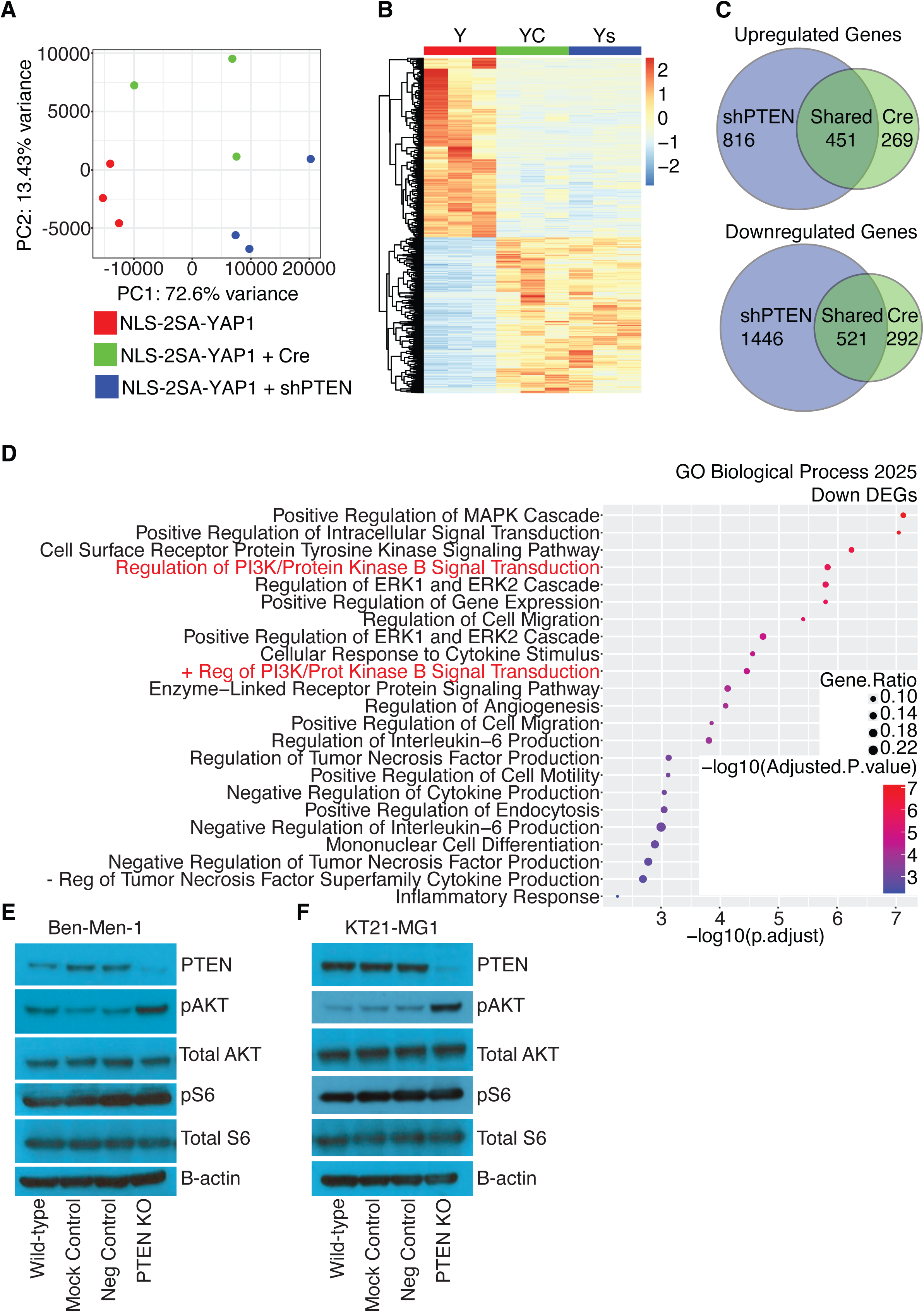
Bulk RNA-seq from mouse models distinguishes *Pten* status *in vivo*. Principal component analysis (PCA) of bulk RNA-seq data from mouse tumors (**A**). Heatmap of differentially expressed genes across mouse tumor models (**B**). Venn diagrams of upregulated and downregulated differentially expressed genes in *Pten*-deficient mouse tumors relative to 2SA-YAP1–driven tumors (**C**). Bubble plot of shared Gene Ontology (GO) Biological Process terms derived from cross-species analysis (**D**). Western blot analysis of PTEN and downstream signaling components in Ben-Men-1 and KT21-MG1 human meningioma cells following CRISPR interference–mediated PTEN knockdown (**E-F**).

To analyze the presence of these signatures in human meningiomas, we first converted the differentially expressed mouse genes to their human orthologs. Then, we identified differentially expressed genes in the human meningioma samples with concurrent chromosome 22q and 10q loss in the aggressive NF2 mutant cluster compared to benign NF2 mutant samples, and performed gene ontology (GO) analysis on these samples (Suppl. Table S4D). Finally, we assessed the overlap between mouse and human datasets (Suppl. Fig. S4B-C).

We next performed GO analysis using the set of genes shared between the two *Pten*-deficient models against the constitutively active YAP1 model (Suppl. Table S4E). Gene set enrichment showed reduced enrichment scores for several GO Biological Process gene sets in *Pten*-deficient tumors. Among the top 100 significant GO Biological Processes terms across species, almost a quarter (23) were shared between mouse and human datasets. These 23 shared GO gene sets were visualized for the mouse tumors, revealing that *Pten*-deficient tumors exhibited downregulation of multiple canonical oncogenic pathways, including PI3K and MAPK signaling (Fig. 4D).

Since PTEN loss did not produce concordant increases in both oncogenic transcription and pathway activity, we sought to define the role of PTEN at the protein level in human NF2 mutant, PTEN wild-type meningioma cell lines, Ben-Men-1 (grade 1) and KT21-MG1 (grade 3) (Suppl. S4D). To do so, we used CRISPR interference gene editing technology to generate *in vitro* PTEN knockouts using these cell lines. Compared with wild-type, mock-transfected controls (transfection without Cas nuclease or guide RNA), and CD8A-targeting negative controls (CD8A is universally not expressed in these lines), PTEN knockout in Ben-Men-1 cells substantially reduced PTEN protein levels and increased AKT phosphorylation, indicating PI3K/AKT/mTOR pathway activation (Fig. 4E). KT21-MG1 cells showed a similar increase in AKT phosphorylation upon PTEN loss (Fig. 4F).

With the knowledge that bulk RNA-seq inherently measures average gene expression across all populations of cells within a sample, coupled with the validation that PTEN knockout increases PI3K signaling in human meningioma cells, we wanted to investigate whether the downregulation of canonical oncogenic signaling in bulk RNA-seq data reflected complex feedback mechanisms or other specific microenvironmental contributions. To resolve this, we turned to single-cell approaches.

### Single-cell RNA-seq from human meningiomas recapitulates *Pten*-deficient mouse model biological signatures

To determine whether the downregulation of PI3K/AKT/mTOR/S6 axis-related GSVA in our *Pten*-deficient mouse models and bulk human RNA-seq data reflected tumor cell-intrinsic effects rather than non-tumor cell contributions, we analyzed single-cell RNA-seq data from human meningiomas (*11*). First, we compiled a set of 49 immune cell marker genes to distinguish neoplastic from non-neoplastic populations (Suppl. Table S5A). Then, we generated a single-cell UMAP (Fig. 5A). After excluding immune cells, we obtained a tumor cell-enriched map, and colored it by WHO Grade and the Heidelberg Methylation Classifier (Fig. 5B-C). Consistent with the bulk RNA-seq data, malignant cells—characterized by numerous CNAs including focal homozygous deletions on chromosome 9p at the CDKN2A/B locus—exhibited significantly lower PTEN expression than Ben-1 type cells—characterized by isolated chromosome 22q loss (Fig. 5D).

**Fig. 5.** Single-cell RNA-seq from human meningiomas recapitulates *Pten*-deficient mouse model biological signatures. UMAP of all cells from human meningioma single-cell RNA-seq datasets (**A**). Tumor cell–enriched UMAP after exclusion of immune populations, colored by WHO grade and Heidelberg methylation class (**B-C**). PTEN expression across benign and malignant tumor cell populations (**D**). Bulk RNA-seq UMAP colored by chromosome 22q and chromosome 10q status (**E**). Copy-number alterations overlaid on the tumor cell UMAP, highlighting chromosome 22q loss and chromosome 10q loss (**F-G**). Comparison of PI3K-related GSVA scores across tumor cell clusters with isolated chromosome 22q loss versus concurrent chromosome 22q and 10q loss (**H**). Statistical analysis was done with a two-sample t-test, where p > 0.05 (ns), p ≤ 0.05 (*), p ≤ 0.01 **(**)**, p ≤ 0.001 (***), and p ≤ 0.0001 (****), unless otherwise indicated.

We next overlaid inferred CNAs onto the tumor cell UMAP. Chromosome 22q loss, the hallmark alteration of NF2 mutant meningiomas, which was located in the top half of the bulk-RNA seq UMAP, was present in nearly all tumor cells, with only a few clusters retaining intact 22q (Fig. 5E-F). The tumor cell UMAP was colored for chromosome 10q status, and these genomic profiles corresponded with high WHO grade tumors and the malignant subtype defined by the Heidelberg methylation classification system (Fig. 5G). Additional recurrently altered chromosome arms, including 1p, 6q, and 14q, were also mapped on the tumor cell UMAP, compared to their bulk counterparts, and revealed that malignant samples frequently harbor multiple arm-level deletions (Suppl. Fig. S5A-F).

Cells with concurrent loss of chromosomes 22q and 10q exhibited significantly lower PI3K-related GO Biological Process scores than clusters with isolated chromosome 22q loss (Fig. 5H, Fig. S5G). These findings suggest that tumor cells recapitulate the PI3K pathway downregulation observed in bulk analyses of aggressive mouse and human tumors. Together, the data support a model in which complex cellular feedback mechanisms suppress PI3K pathway gene expression following PTEN loss, revealing a cell-intrinsic adaptation in aggressive NF2 mutant meningiomas.

### *Pten*-deficient mouse models recapitulate aggressive NF2mut human meningioma biology

To assess the concordance between our mouse models and human tumors, we defined custom gene sets from genes significantly upregulated and downregulated in *Pten*-deficient mouse models and applied GSVA. This approach enabled us to move beyond standard GO terms and directly assess how disease-specific gene signatures shape broader biological problems.

GSVA scores for the two gene sets, and the ratio between them, were projected onto the human UMAP. This analysis revealed that Cre-mediated *Pten* knockout genes highlighted the aggressive NF2 mutant regions of the UMAP (Fig. 6A). Kaplan-Meier analysis showed that patients in the upper quartile of these GSVA scores experienced shorter times to recurrence, an effect that was more pronounced when stratifying by chromosome 10q status (Fig. 6B).

**Fig. 6.** *Pten*-deficient mouse models recapitulate aggressive NF2mut human meningioma biology. UMAP of human bulk RNA-seq colored by the ratio of GSVA scores for Cre-mediated *Pten* knockout gene sets, and corresponding Kaplan-Meier curve stratified by chromosome 10q status (**A-B**). UMAP of human bulk RNA-seq colored by the ratio of GSVA scores for shRNA-mediated *Pten* knockdown gene sets, and corresponding Kaplan-Meier curve stratified by chromosome 10q status (**C-D**). UMAP of bulk RNA-seq colored by the ratio of GSVA scores for Pten-deficient gene sets shared amongst the two *Pten*-deficient tumors, and corresponding Kaplan-Meier curve stratified by chromosome 10q status (**E–F**). UMAP colored by highly aggressive, intermediate, and benign NF2 mutant and NF2 wild-type clusters and its corresponding Kaplan-Meier curve, where Ben NF2mut vs. NF2wt p = 0.0232, Ben NF2mut vs. Int NF2mut p = 1.2E-8, Ben NF2mut vs. Highly Agg NF2mut p = 2E-16, NF2wt vs. Int NF2mut = 0.0015, NF2wt vs. Agg NF2mut = 2E-16, Int NF2mut vs. Highly Aggressive NF2mut = 9.0E-16 (**G-H**). GO analysis of shared mouse and human genes (**I**). Statistical analyses were conducted using the log-rank (Mantel–Cox) test. Significance thresholds were defined as p > 0.05 (ns), p ≤ 0.05 (*), p ≤ 0.01 **(**)**, p ≤ 0.001 (***), and p ≤ 0.0001 (****), unless otherwise indicated.

shRNA-mediated *Pten* knockdown showed comparable patterns, and recurrence analysis indicated that shRNA-mediated *Pten* knockdown was even more predictive of poor human outcomes than Cre-mediated *Pten* knockout, supporting earlier observations that shRNA-mediated *Pten* loss was more aggressive than Cre-mediated *Pten* loss *in vivo* (Fig. 6C-D).

Combining transcriptional profiles of Cre- and shRNA-mediated *Pten* loss further enhanced the predictive power for human outcomes (Fig. 6E–F). Maps colored by up- or downregulated genes alone also showed predictive patterns, although they were less robust than the ratio-based analysis (Suppl. Fig. S6A–G).

Samples with high GSVA scores were enriched in the aggressive NF2 mutant region of the UMAP and had very poor clinical outcomes. This finding mirrors other studies that have used the UCSF 34-gene signature to highlight this highly aggressive subcluster and show that this region of the map has a large population of patients with poor outcomes. Kaplan-Meier analysis supported these distinctions: patients with highly aggressive NF2 mutant tumors recurred earliest, followed by those with intermediate NF2 mutant, NF2 wild-type, and benign NF2 mutant tumors (Fig. 6G-H). Identifying genes shared between *Pten*-deficient mouse models and humans revealed downregulation of PI3K-associated GO Biological Process gene sets (Fig. 6I, Suppl. Table S6A). Using these shared genes, finding the ratio of up- and downregulated genes, and coloring the map again highlighted the aggressive NF2 mutant region of the UMAP (Suppl. Fig. S6H).

## DISCUSSION

While benign NF2 mutant meningiomas exhibit few genomic alterations beyond chromosome 22q loss, aggressive NF2 mutant tumors display extensive arm-level chromosomal aberrations. We focused on chromosome 10q, which harbors the well-characterized tumor suppressor PTEN. We show that in human patients, loss of chromosome 10q and PTEN is associated with significantly shorter times to recurrence. Using our previously established genetically engineered mouse models, we demonstrated that *Pten* loss significantly enhanced the malignancy of YAP1-driven models. These data mirror findings from glioblastoma, where PTEN loss (frequently mediated via chromosome 10q monosomy and mutational inactivation of the remaining PTEN allele) represents a central oncogenic mechanism.

While both shRNA-mediated *Pten* knockdown and Cre-mediated *Pten* knockout showed a similar trend of accelerated tumor formation in YAP1-driven meningiomas, shRNA-mediated *Pten* knockdown was significantly more aggressive. Cre-mediated *Pten* knockout was a non-significant trend, which may suggest that complete loss of *Pten* is somehow detrimental to the cancer cells, and they might need a residual amount of *Pten* to drive highly aggressive tumorigenesis.

In meningiomas, PTEN loss activates PI3K signaling at the protein level, but paradoxically suppresses transcripts downstream of the PI3K pathway in both mice and humans. Since PTEN loss did not lead to concordant activation of PI3K pathway components at the RNA level, this phenomenon may reflect cell type-specific feedback mechanisms, the basis of which remains unclear and warrants further investigation.

Integrating our mouse meningioma data with large human meningioma datasets strengthens the confidence in and clinical relevance of our conclusions. The gene expression program induced by *Pten* loss in mice is observed not only in aggressive, PTEN-deficient human meningiomas, but also in aggressive meningiomas that retain intact PTEN. This suggests that while PTEN loss is one route to malignancy, additional mechanisms can produce similar transcriptional signatures and biological phenotypes.

Using RCAS/tv-a technology and other genetically engineered mouse models, we can now directly test causality. For example, through conditional knockout or shRNA-mediated knockdown of candidate tumor suppressors on recurrently lost chromosomes, such as chromosome 1, 6, and 14, we can now assess effects on tumor latency, biology, and therapeutic response. Importantly, since these models faithfully recapitulate aggressive human disease, they offer a tractable *in vivo* platform to study meningioma progression and evaluate novel therapeutic strategies and resistance mechanisms for clinically challenging tumors.

In the future, we envision mapping patients’ transcriptomic profiles onto this reference landscape in real time, enabling clinicians to compare individual tumors with their patients’ “nearest neighbors.” This approach could have significant diagnostic and prognostic value, particularly for meningiomas with chromosome 10q loss and other CNAs.

A particularly powerful aspect of this pipeline is its broad applicability across disease contexts. Transcriptional profiling of preclinical models in mice and other model organisms can be directly compared with that of humans, enabling rigorous translational insight and intervention. Moreover, this framework is readily scalable and extends beyond transcriptomics: dimension-reduced landscapes can integrate genomic, metabolomic, and proteomic data from both preclinical and clinical cohorts, supporting multi-omic, cross-species analyses through joint dimensionality reduction and integrative modeling. By embedding human disease biology as a benchmark for model fidelity, this approach reshapes how preclinical models are generated and validated, strengthening the robustness, reproducibility, and interpretability of biological mechanisms and therapeutic opportunities.

## MATERIALS AND METHODS

### Study Design

This study leveraged both human meningioma samples and genetically engineered mouse models to establish a causal role for PTEN loss in driving aggressive NF2 mutant tumors. Human bulk and single-cell RNA-seq datasets were analyzed to define transcriptional subtypes, identify chromosomal alterations, and correlate these features with clinical outcomes. Complementary preclinical models included primary arachnoid cell cultures and RCAS/tv-a tumors engineered to express constitutively active YAP1 alongside conditional or shRNA-mediated *Pten* loss, in both *Cdkn2a* wild-type and knockout backgrounds. Biological signatures were assessed using histology, immunohistochemistry, and RNA-seq. Finally, cross-species projection of mouse gene signatures onto human datasets validated the preclinical models and directly linked PTEN loss to molecular and phenotypic hallmarks of tumor aggressiveness, providing mechanistic evidence of causality.

### Bulk RNA Sequencing Data from Human and Mouse Meningiomas

Bulk RNA sequencing data from human meningiomas have been previously published (*10, 11*). For analysis and visualization of human meningioma tumors, normalized and raw bulk RNA-seq counts, UMAP coordinates, genomic alterations and clinical metadata for 1,298 human meningioma tumors were used.

Extracranial mouse tumors were flash-frozen in liquid nitrogen for *in vivo* RNA expression analysis. RNA extraction, library preparation, and poly(A)-capture sequencing was performed by the Genomics and Bioinformatics Shared Resource at Fred Hutch Cancer Center. RNA-seq reads were aligned to the UCSC mm10 using STAR and counted for gene associations against the UCSC genes database with HTSeq.

The code used to process and analyze the data is available at https://github.com/sonali-bioc/Parrish_PTEN_in_meningioma_Manuscript/.

This research was supported by the Genomics & Bioinformatics Shared Resource, RRID:SCR_022606, of the Fred Hutch/University of Washington/Seattle Children’s Cancer Consortium (P30 CA015704).

### Single-Cell RNA Sequencing from Human Meningiomas

Single-cell RNA-seq data of human meningioma was obtained from the data repository of the Department of Neuropathology at the University Hospital Heidelberg, and is available upon request.

Seurat objects were constructed for 26 single-cell RNA-Seq samples of human meningiomas. Cells from the 26 single-cell RNA-Seq samples were further filtered to remove immune cells based on 49 marker genes spanning a variety of immune cell types (See Suppl. Table S5A). Next SCTransform from Seurat was used to process the data, followed by building a UMAP and forming clusters. Metadata extracted for the remainder cells was used to divide cells into different groups for CNS WHO grade and methylation cluster (MC) status. Dotplots and violin plots for key genes were made using the function DotPlot() and VlnPlot() respectively from Seurat. FindMarkers() from Seurat was used to find differentially expressed genes between the different MC status (ben-1 vs ben-2, ben-1 vs int-A, ben-1 vs int-B, ben-1 vs mal). R package, R function, DoHeatmap() were used to make heatmaps. All analysis was done using R 4.3.3 and Seurat (v 5.0.0).

The code used to process and analyze the data is available at https://github.com/sonali-bioc/Parrish_PTEN_in_meningioma_Manuscript/.

### Kaplan-Meier Curves

Patient samples were divided into groups based on the expression of the gene of interest; high expression (higher than 3^rd^ quartile) vs low expression (lower than 1^st^ quartile). Kaplan-Meier curves were generated using the information on time to recurrence of each sample. To perform the calculations, we only selected samples with known recurrence status (recurrence = yes/no) and known time to recurrence or last follow up. For tumors that were confirmed as non-recurrent however with no last follow up date, a default of 315 months was used as “Months of No Recurrence.” Kaplan-Meier curves were plotted using the R packages “ggsurvfit” and “survminer.”

### Differential Gene Expression Analysis

R package DESeq2 was used to find differentially expressed genes (DEGs) between mouse models and clusters. A threshold of fold change of 1.5 (or logFC of 0.58) and a p-adjusted value < 0.05 was used to determine significantly regulated DEGs. Upregulated and downregulated genes in each cluster are listed in Suppl. Table S4A-D. Gene sets were made using upregulated and downregulated genes in *Pten*-deficient mice relative to *Pten* wild-type mice. Shared DEGs for *Pten* knockdown and *Pten* knockout models were identified independently, and the overlapping gene set was subsequently determined. Mouse genes were mapped to their corresponding human orthologs using BioMart to link the respective Ensembl databases.

### Pathway Analysis

Statistically significant upregulated or downregulated genes were analyzed using Enrichr to determine the enriched biological signature (*19*). Gene Ontology Biological Processes (GO BP) terms that were statistically significant (adjusted *p* value <0.05) were selected and dot plots were generated using ggplot2 (top 23 GO terms included). Complete lists of GO terms were attached as supplemental information.

### Generation and Transduction of Murine Arachnoid Cells

DF-1 cells were transfected using the procedure outlined in “Transfection of RCAS Viruses.” Viral supernatant from transfected DF-1 cells was filtered through a 0.45 µm filter. Murine arachnoid tissue was dissected from the ventral side of the skull of (*N*)/tv-a; *Cdkn2a*^−/−^; *Pten*^fl/fl^ mice and dissociated in phosphate-buffered saline (PBS), followed by mechanical dissociation using syringes and needles. Arachnoid cells were isolated, collected as a pellet, and resuspended in the viral supernatant from transfected DF-1 cells in a 24-well flat-bottom plate. The arachnoid cells were gradually expanded and continuously transfected with viral supernatant from DF-1 cells for 7-10 days.

### Cell Culture

Chicken fibroblast DF-1 and murine arachnoid cells were maintained in Dulbecco’s Modified Eagle Medium (DMEM, REF: 11995-065) supplemented with 10% fetal bovine serum (FBS) and 1% penicillin-streptomycin. Ben-Men-1 cells were obtained from DSMZ (ACC 599), while CH157-MN and KT21-MG1 cells were kindly gifted by Dr. Hiroaki Wakimoto (Brain Tumor Research Center, Massachusetts General Hospital) and Dr. Jonathan Chernoff (Fox Chase Cancer Center), respectively. CH157-MN and KT21-MG1 were cultured in DMEM supplemented with 10% FBS and BM1 were maintained in DMEM supplemented with 20% FBS and 1% penicillin-streptomycin. All mammalian cells were incubated in a humidified incubator at 37°C with 5% CO_2_. Avian cells were incubated in a humidified incubator at 39 °C with 5% CO_2_.

### Cas9:sgRNA RNP Nucleofection

Two synthetic single guide RNAs (sgRNAs) targeting PTEN, each containing 2’-O-methyl phosphorothioate modifications, were purchased from Synthego. The sgRNA sequences were 5’-AAAAGGAUAUUGUGCAACUG-3’ and 5’-UGUGCAUAUUUAUUACAUCG-3’. Each sgRNA was reconstituted to 100 pmol/µL in nuclease-free 1X TE buffer (Tris-EDTA, pH 8.0). The sNLS-SpCas9-sNLS nuclease (Aldevron, #9212) was diluted from 61 pmol/µL to 10.17 pmol/µL. Complete Nucleofector Solution was prepared by combining the Nucleofector Solution and the supplied Supplement at a 4.5:1 ratio. Ribonucleoprotein (RNP) complexes were assembled by first adding reconstituted sgRNAs to the Complete Nucleofector Solution, followed by the diluted Cas9, for a final volume of 20 µL using a Cas9:sgRNA ratio of 1:2. The RNPs were incubated at room temperature for 15 minutes and then placed on ice until use. A total of 2.0 × 10⁵ cells were harvested, washed with PBS, and resuspended in 20 µL of the RNP mixture. Cells were electroporated using the Amaxa 96-well Shuttle System (program EN-138), recovered in pre-warmed media, and then plated dropwise into 12-well plates. Media was changed 24 hours post-nucleofection.

### Western Blotting

Protein lysates were collected using RIPA buffer supplemented with 100X EDTA Solution (REF: 1861274) and 100X Halt Protease and Phosphatase Inhibitor Cocktail (REF: 1861281). Protein concentration was quantified using the Bicinchoninic Acid (BCA) assay (REF: PC201323), with bovine serum albumin (BSA) as a standard. Equal amounts of protein were denatured with 4X Invitrogen NuPAGE LDS Sample Buffer (REF: NP0007) and heated at 95°C for 10 minutes. Samples were loaded onto a 4-12% Bis-Tris polyacrylamide gel (REF: NP0323BOX) for electrophoresis and run at 100 V for 2 hours with MOPS SDS Running Buffer (REF: NP0001). After separation, proteins were transferred to an Immun-Blot PVDF membrane (REF: 1620177) at 20 V for 1 hour with NuPAGE Transfer Buffer (REF: NP00061). Membranes were blocked with 2% BSA in Tris-buffered saline with 0.1% Tween-20 (TBS-T) and incubated overnight at 4°C with primary antibodies diluted in 2% BSA in TBS-T. Following washing with TBS-T, appropriate secondary antibodies were applied, also diluted in 2% BSA in TBS-T. Membranes were then developed on ProSingle Blotting Film (REF: 30-810L) using Amersham ECL Western Blotting Detection Reagents (REF: RPN2106) and imaged using a designated imaging system.

Blotting
| Antibody | Supplier | Catalog # |
| --- | --- | --- |
| B-actin | Sigma | A1978 |
| HA-Tag | Cell Signaling Technology | 2367 |
| HA-Tag | Cell Signaling Technology | 3724 |
| Phospho-AKT | Cell Signaling Technology | 4060 |
| Phospho-S6 | Cell Signaling Technology | 4858 |
| PTEN | Cell Signaling Technology | 9559 |
| Total AKT | Cell Signaling Technology | 9272 |
| Total S6 | Cell Signaling Technology | 2217 |

### Generation of RCAS Mouse Meningiomas

Animal experiments were performed in compliance with the Institutional Animal Care and Use Committees (IACUC) at Fred Hutchinson Cancer Center (protocol no. 50842) and adhered to the National Institutes of Health Guidelines for animal welfare. DF-1 cells were transfected with the appropriate RCAS plasmids using X-tremeGENE 9 DNA Transfection Reagent (REF: 06365809001), according to the manufacturer’s protocol. RCAS/tv-a technology was employed to model meningiomas. (*N*)/tv-a; *Cdkn2a*^−/−^; *Pten*^fl/fl^ and (*N*)/tv-a; *Cdkn2a^+/+^* were used for these studies. A 1 µL injection (or 1:1 mixture for combined injections) of a 1 × 10^5^ transfected DF-1 cell suspension was delivered into the subarachnoid space of neonatal (postnatal day 0-3) mice. Mice were monitored for symptoms of tumor burden, including visible extracranial tumors, lethargy, poor grooming, hunched posture, weight loss, and emaciation, or until a predetermined study endpoint (in most cases, 150 days post-injection).

### Incucyte Live-Cell Imaging

Cells were seeded at 5 × 10^2^ cells per well in a tissue culture-treated 96-well plate (REF: 353072) containing DMEM supplemented with 10% FBS. Live-cell imaging was performed using the Incucyte ZOOM system (Essen) every 24 hours for 5 days. Phase object confluency, normalized to Day 0, was used for analysis. This research was supported by the Cellular Imaging Shared Resource RRID:SCR_022609 of the Fred Hutch/University of Washington/Seattle Children’s Cancer Consortium (P30 CA015704).

### Histology

For routine histological examination, mouse brains and peripheral tumors were formalin-fixed, paraffin-embedded, decalcified in formical-4 (if tissue contained bone), sectioned, and stained with hematoxylin and eosin (H&E). Immunohistochemistry (IHC) was performed on the Roche Ventana DISCOVERY ULTRA. Option consisted of 10% Normal Goat Serum in 2% bovine serum albumin (BSA) and detection consisted of biotinylated goat anti-rabbit (1:300) in 2% BSA in PBS. Antibodies were also diluted in 2% BSA in PBS. A list of antibodies used is provided in “Western Blotting” section.

### Immunofluorescence

Deparaffinization and antigen retrieval protocols were performed. Slides were rehydrated through graded alcohols to water and permeabilized in 0.3% Triton-X in 1X PBS for 40 minutes on a rocker. Slides were blocked for 1 hour at room temperature in 2% BSA in PBS supplemented with 5% donkey serum and 0.1% Triton X. Primary antibodies were diluted in 2% BSA in PBS and incubated overnight at 4°C: rat anti-HA (1:100) and rabbit anti-PTEN (1:200). Slides were washed in PBS and incubated for 1 hour at room temperature with species-appropriate anti-rat Alexa Fluor 488 and anti-rabbit Cy3. Nuclei were counterstained with Hoechst, slides were rinsed, coverslipped with 70% glycerol, and stored long-term at 4°C.

### Statistical Analyses

Statistical analyses were performed using GraphPad Prism 10 and R. For comparisons of continuous variables across multiple groups, a one-way analysis of variance (ANOVA) was used. Kaplan-Meier survival curves were generated to assess recurrence-free survival, and differences between groups were evaluated using the Log-rank (Mantel-Cox) test. Gene expression differences in bulk RNA-seq and single-cell RNA-seq datasets were evaluated using DESeq2. For gene set variation analysis (GSVA), enrichment scores were compared using one-way ANOVA. Significance thresholds were defined as: p > 0.05, not significant (ns); p ≤ 0.05 (*); p ≤ 0.01 (**); p ≤ 0.001 (***); p ≤ 0.0001 (****). Exact p-values are reported in the figure legends or main text where applicable.

## Acknowledgements

We would like to thank Debra Kumasaka, Kelly Grissom, and Linda Lew for administrative assistance and support throughout these experiments and the publication process.

## Author Contributions

Conceptualization: AGP, FS, ECH

Methodology: AGP, HNT, SA, CHM, PS, FS

Investigation: AGP, HNT, SA, CHM, PS, FS

Visualization: AGP, HNT, SA

Funding acquisition: ECH

Project administration: ECH

Supervision: FS and ECH

Writing – original draft: AGP, FS, ECH

Writing – review & editing: AGP, FS, ECH

## Data and materials availability

All data are available in the main text or the supplementary materials and are also available through the corresponding author upon reasonable request.

