## Supplementary figures and images for "PTEN deficiency linked to chromosome 10q loss leads to aggressive NF2 mutant meningioma biology"

### Supplemental Figures

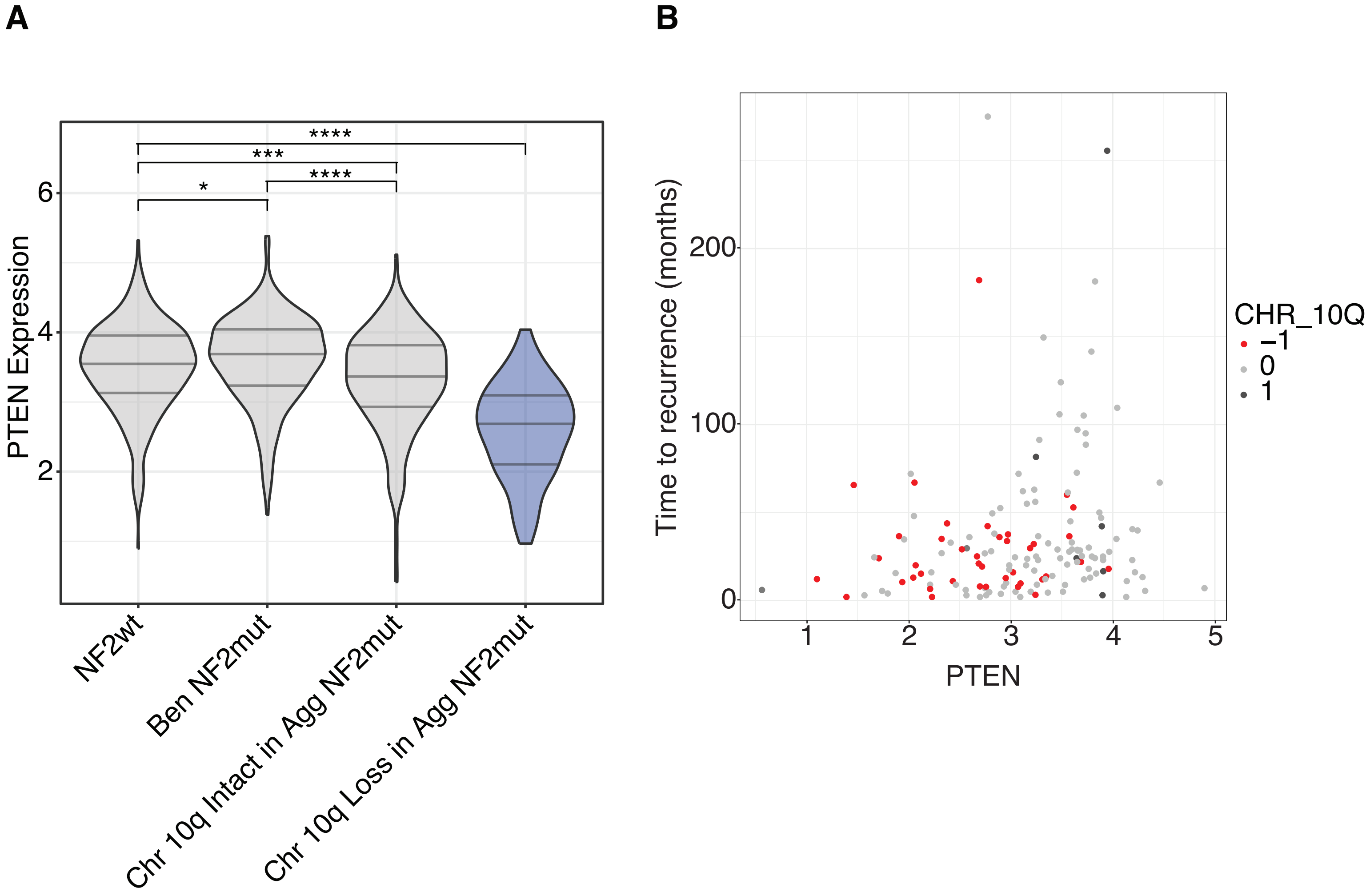

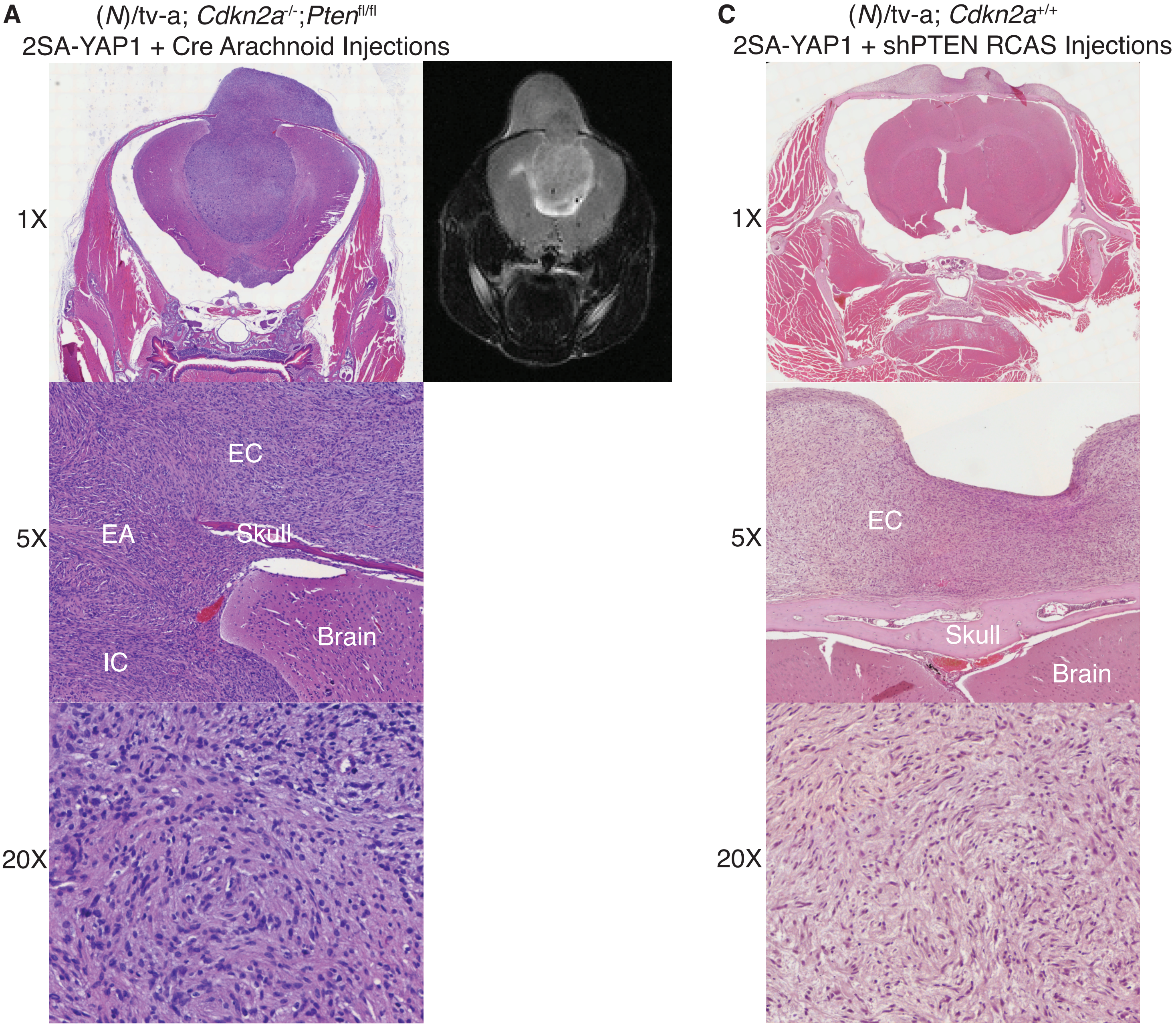

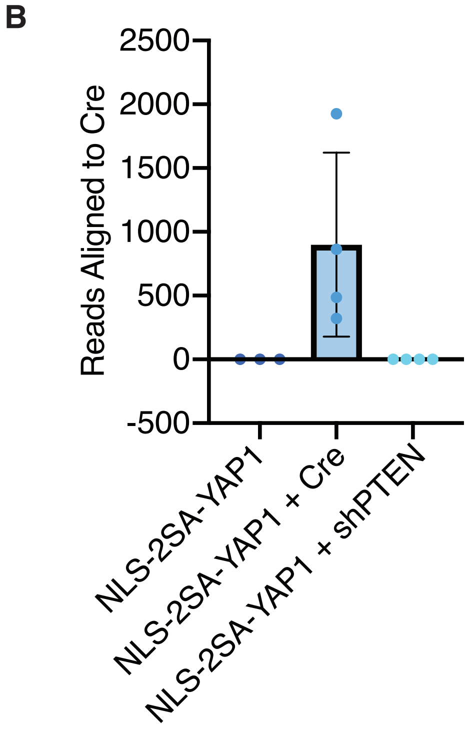

**Suppl. Fig. 4.**

**A**

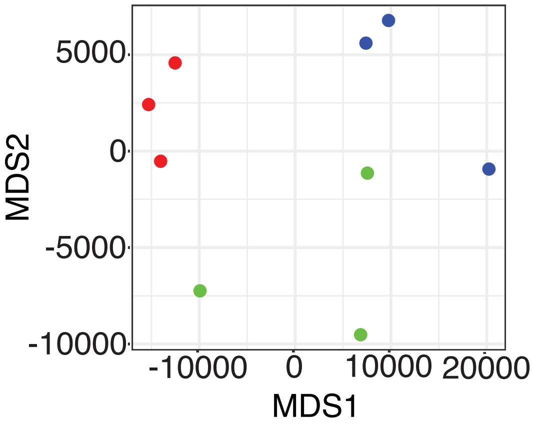


**B**

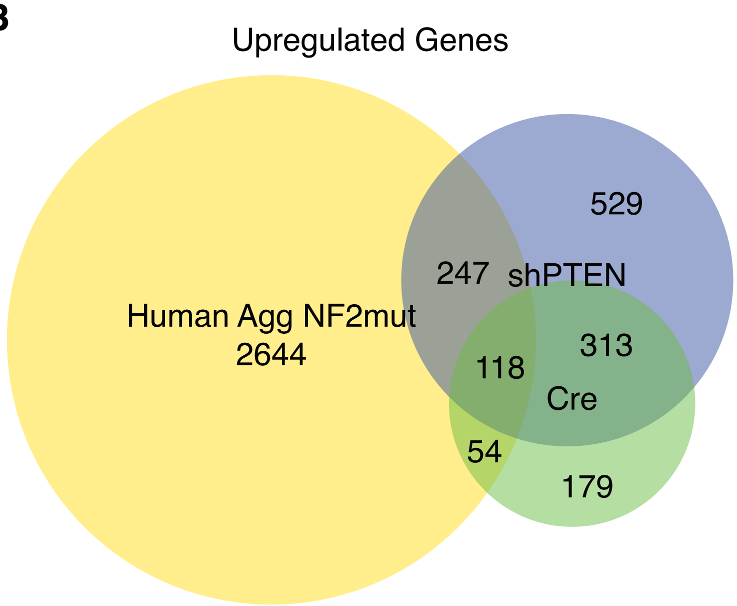


**C**

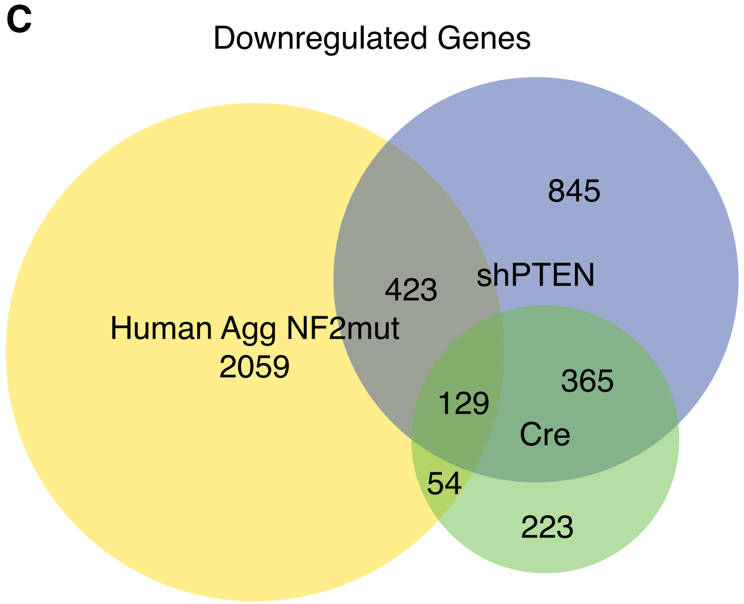


**D**

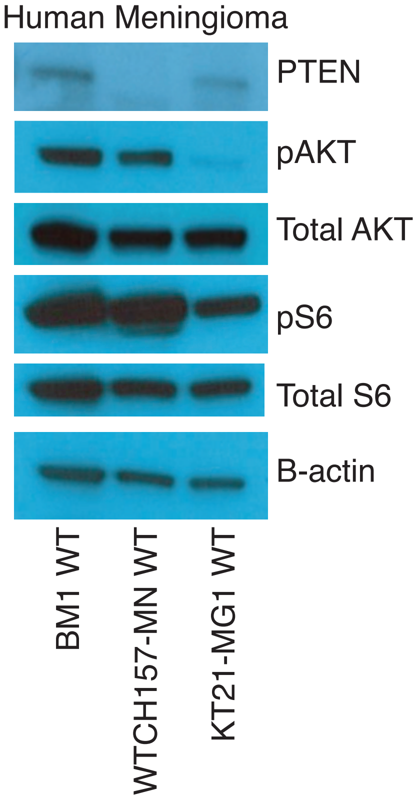

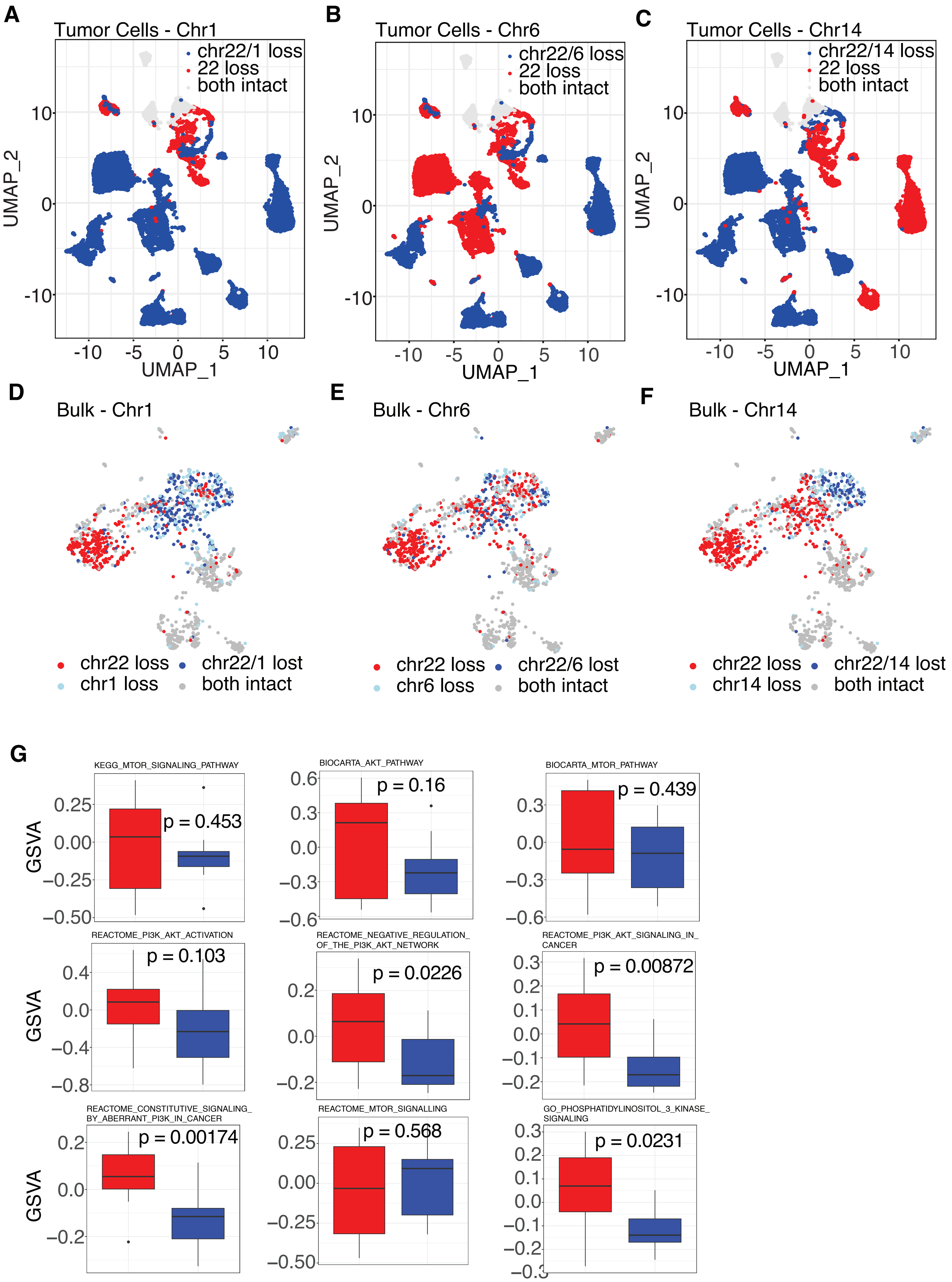

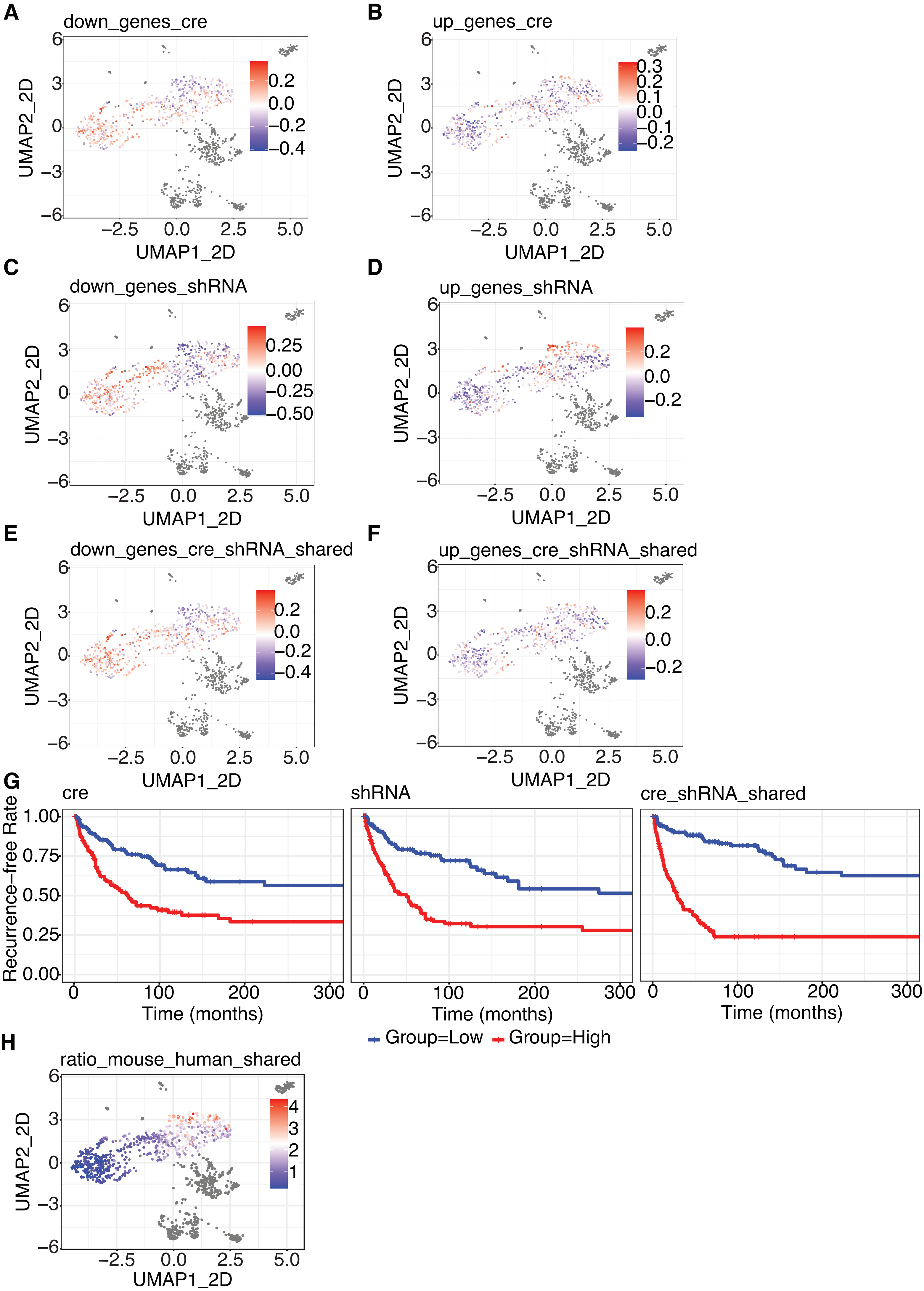
